# Differential Survival In Carrion Crows Through Eavesdropping on Human Speech: An Agent-Based Model

**DOI:** 10.1101/2023.09.15.557945

**Authors:** Sabrina Schalz

**Affiliations:** Middlesex University, Department of Psychology, London, UK

**Keywords:** Carrion crow, Heterospecific communication, Eavesdropping, Agent-based model

## Abstract

Wild carrion crows eavesdrop on human speech and respond more strongly to it than to the calls of pigeons or parakeets. However, it remains unclear whether this behaviour provides any benefit to them, and if so, under which environmental conditions such a benefit could be expected. I used an agent-based model to examine the effect of eavesdropping on the survival of a carrion crow under different environmental conditions. The crow could gain points from foraging, but also had to avoid encounters with a human in order to not lose points. In addition to eavesdropping, it could visually scan the patch for human presence and, once detected, could keep track of the human’s location. I found that eavesdropping led to higher final scores and higher rates of survival than visual scanning alone if the human density on the patch was elevated, or if there were visual obstacles reducing the accuracy of visual scans. Such environmental conditions are particularly common for urban crows foraging in areas such as public parks. The model provides a proof of concept for future empirical research into the potentially differential survival of urban crows eavesdropping on speech.

## 1. Introduction

### 1.1 Eavesdropping on Speech

One environmental condition that all urban populations have to account for is human presence. Foraging in an urban habitat may require individuals to adjust their behaviour to direct and indirect human disturbance (both acute disturbance such as humans walking close-by, as well as long-term disturbance such as habitat degradation), while also exploiting benefits resulting from human presence, including access to anthropogenic food sources, both incidental (from a rubbish bin) and intentional (from a bird feeder). Frid and Dill (2002) suggested that “disturbance stimuli should be analogous to predation risk” and list vigilance as a response to disturbance. These disturbance stimuli include vocalizations of other species, as eavesdropping on heterospecific vocalizations allows individuals to detect the presence of a predator (Magrath et al., 2015).Common ravens (*Corvus corax*), for instance, respond to playback of jackdaw (*Corvus monedula*), blue jay (*Cyanocitta cristata*), European jay (*Garrulus glandarius*), and laughing gull (*Leucophaeus atricilla*) alarm calls with flight or vigilance (Davídková et al., 2020; Nácarová et al., 2018).

In some species, this perception of heterospecific vocalizations extends to human speech as well. Mountain lions (*Puma concolor*), as well bobcats (*Lynx rufus*), striped skunks (*Mephitis mephitis*) and Virginia opossums (*Didelphis virginiana*) in the Santa Cruz Mountains showed reduced activity and avoided areas where recordings of human speech were played (Suraci et al., 2019). Both captive carrion crows (*Corvus corone*) as well as wild Western Australian magpies (*Gymnorhina tibicen dorsalis*) respond more to unfamiliar than to familiar human voices, which may be due to the unknown level of threat posed by an unfamiliar person (Dutour et al., 2021; Wascher et al., 2012). As some people attempt to scare or chase crows away (Clucas & Marzluff, 2012; Schalz, 2021) and crows are sometimes killed by humans (Animal Aid, 2017; Fujioka, 2020; The Royal Parks, 2018), humans may be perceived as a threat by wild crows.

Playback experiments with human speech have shown that wild carrion crows respond more to speech than they do to playback of parakeet or pigeon calls (Schalz, 2023), responding significantly more often with flight behaviours when hearing speech. Other species, such as large-billed crows (*Corvus macrorhynchos*) (Schalz & Izawa, 2020) or domestic dogs (*Canis familiaris*) (Cuaya et al., 2022; Mallikarjun et al., 2022) respond differently to two different human languages following natural exposure, without needing explicit training or encouragement. Eavesdropping on speech could therefore be beneficial to wild crows by reducing the risk of coming too close to a potentially threatening human. This may be particularly beneficial in urban areas where human density is high and where crows often forage in areas such as public parks. However, eavesdropping is also less precise than visual detection. Broadcasted acoustic signals only provide moderate locatability (Krebs & Davies, 1987), which is enough to trigger a vigilance response but less useful to locate the source.

This lower precision could potentially result in false alarms when the human is present but far away. Such a false alarm would be costly as it could lead the crow to abandon the foraging opportunity unnecessarily. Not responding to cues to human presence can also be costly in the form of injury from hostile encounters (referred to predation throughout the paper), as some people will chase after crows (Schalz, 2021).

### 1.2 Modelling Approaches

Whether a vigilance behaviour such as speech eavesdropping actually leads to differential survival is challenging to ascertain in the field, but can instead be modelled theoretically. For instance, Brown (1999) used a mathematical model to study trade-offs between vigilance and foraging patch use. In this model, the baseline foraging rate was negatively correlated with the vigilance level. The amount of time spent on a patch was reflected in the net rate of energy gain from food intake, as well as the energetic cost of foraging (including vigilance, search and handling costs). The forager’s fitness increased if the net energy gain was positive, and decreased if it was negative. The risk of instant death was constant for a patch and not influenced by the time spent on it. An increased vigilance rate could compensate for low vigilance effectiveness. A higher encounter rate with predators increased the predation risk, whereas higher vigilance rates and odds of escaping the predator without vigilance decreased the predation risk. Overall, survival chances were proportionate to the predation risk and employed vigilance on a given patch.

Agent-Based Models (ABMs) are useful alternatives to mathematical models and have gained popularity in ecological modelling. They are particularly suitable to model variation in individual behaviour, and are invaluable when fieldwork is overly impractical or costly. An ABM is a simulation to test out theoretical assumptions, such as how different individuals (agents) interact with each other (DeAngelis & Diaz, 2019). This agent “is an entity with a set of parameters and a defined objective that interacts with other entities within the model to achieve this objective” (Murphy et al., 2020), for example predator and prey interactions (Ringelman, 2014) and predator avoidance (Watzek et al., 2021). Agents can be designed with different physical traits, behaviours, and decision-making choices (DeAngelis & Diaz, 2019). They are placed in an environment (such as grid-based patches, similar to chess boards) with spatial and temporal parameters (Murphy et al., 2020), where each agent will have a specified location and specified movement behaviours (see figure 1).

**Figure 1:**
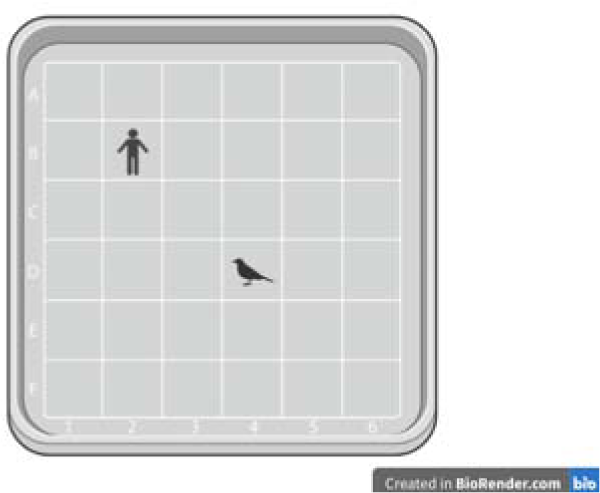
Illustration of the model concept. A crow and a human (two agents) are standing on a patch (the environment) divided into squares. Note that this illustration only shows 6×6 squares for simplicity, but the model was designed with 20×20 squares.

## 2. Methods

The model description follows the ODD (Overview, Design concepts, Details) protocol for describing individual-and agent-based models (Grimm et al., 2006) as updated by Grimm et al. (2020). The model was written in R v.4.0.2 (R Core Team, 2020), and the R script was based on the open-source ABM R script for predator-prey dynamics provided by Duthie (n.d.), with modifications and added behaviours where needed (full script provided as supplementary materials). In addition to the theoretical assumptions outlined by Brown (1999), the model parameters are informed by empirical data of carrion crow ecology. However, as many variable and parameter values used in this model are estimates and will have considerable ranges of variation in real life, the predictive powers of this model are quite limited.

The main aim in this paper is to provide a proof of concept for the fitness benefit that eavesdropping on speech likely provides, and to inspire further research into this area, both empirical by attempting to replicate the predictions made from the model, and theoretical to expand the scope and complexity of the model itself.

### 2.1 Purpose and Patterns

The purpose of the model is to predict under which environmental conditions eavesdropping on human speech provides a benefit to carrion crows, in the form of higher survival odds compared to individuals not using this additional vigilance behaviour. Eavesdropping on speech allows crows to detect human presence, similar to other vigilance behaviours such as visual scanning. The faster and more reliable a crow can detect human presence, the lower the risk of a potentially harmful encounter. The predictions made from the model will guide and inform future empirical research into potentially differential survival of crows eavesdropping on speech in urban areas. In the model, I focused on two central patterns:

*Pattern 1:* The risk of an encounter is relative to human density on a patch, such that higher human densities result in higher risks of encounters. This can be due to either the patch being small, or the number of humans being high, or both. The higher the human density, the quicker human presence must be detected to avoid an encounter.

*Pattern 2:* The risk of an encounter is relative to the accuracy of vigilance behaviours. Crows can visually scan their surroundings and look for humans, but this will be less reliable if there are visual obstacles on the patch. If a human could be standing behind an object (such as a tree, a car etc.), visual scanning could produce false negatives. The more obstacles are on the patch, the lower the accuracy of visual scanning and the higher the risk of false negatives.

### 2.2 Entities, State Variables, and Scales

Two agents, a crow and a human, are placed on a patch divided into 20×20 squares, comparable to a chess board (figure 1). This patch size was chosen to allow encounters to occur regularly, but not constantly. A patch with a discrete size and border (rather than modelling a continuous patch) is analogous to the experimental sites used in (Schalz, 2023), where a grass area (such as a park lawn) was bordered on all sides by trees and/or buildings, and where the crows foraged on the grass but not in the trees. There can thus be several such patches next to each other (such as in a large park) or only one grass patch as modelled here (such as a single football field in a suburban neighbourhood without other lawn areas of comparable size and public access), but each patch still has distinct borders, reflected in the design of the patch in the present model.

Each square on the patch has a size of 1m^2^ and provides food without depletion. Both agents have a state variable for their x-coordinate and one for their y-coordinate, to represent their position on the patch. This is a whole number between 1 and 20 and refers to a specific square on the patch (for example, square 1;5 would be the square that is in the first column on the x-axis and the fifth row on the y-axis).

The model runs through 300 iterations. Each iteration corresponds to 30min in a 7.5h day (based on activity patterns observed during fieldwork (Schalz, 2023) in winter in the UK, where both carrion crows and humans arrived on the patch shortly after the 8am sunrise, and had left by sunset at 4pm). This resulted in 15 iterations in a day spread across a total of 20 days. Within each iteration are 100 time units *t*, and a full run of the model therefore adds up to 30,000 *t*. Splitting an iteration into 100 *t* (corresponding to 18s) was chosen over 60 *t* (corresponding to an even 30s) for the sake of simplicity in calculations. The length of iterations were chosen to create a realistic balance between a crow’s daily energy expenditure relative to energy intake opportunities (i.e. energy costs and foraging, see below), and *t* were chosen to break this down into smaller time steps to allow for a series of actions by the crow and the human. However, the model would be expected to work equally well with different time step sizes.

In each *t*, the agents execute at least one behaviour. When a new iteration begins, the set-up resets and the two agents are again randomly placed on the patch. This nested design of time units within larger iterations was chosen so that behaviours could occur on a fairly granular level, but that the crow would have to start over several times in a day, to reflect natural conditions of wild crows foraging across different patches throughout the day.

In addition to the x and y-coordinate location on the patch, the crow has a state variable for points (so a total of two state variables for the crow, and only one for the human agent). Points are integer values that measure the crow’s health throughout the model, such that 0 points correspond to its death. The points were scaled around calorie values, so that 1 point corresponds to 0.01kcal. Based on formula 1 for daily energy expenditure (DEE) values for passerines (Department for Environment, 2002) and the weight of a carrion crow (Hume, 2018), a crow needs 208kcal per day. This corresponds to 14kcal per iteration, or 1,400 points.

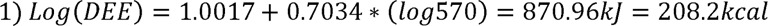

The crow starts the model with 21,000 points (210 kcal). To represent the energy expenditure of staying alive, each iteration costs 1,400 points. If the crow does not have enough points to enter another iteration (i.e. it has a total score of less than 1,400 points), it dies and the model ends. This is to reflect the crow’s ongoing energy requirements to maintain life and body functioning. Each iteration, the crow then gains and loses points by executing predefined behaviours (see 2.3). These gains and costs were determined individually for each behaviour based on empirical data and preliminary tests of the model. Starting the model with 21,000 points corresponds to a day’s worth of calories, to allow some room for potential point losses in early iterations (rather than starting with 0 or near 0 points, where the crow would die after just one encounter with a human, see 2.3).

### 2.3 Process Overview and Scheduling

The model consists of several behaviours executed by the agents (see figure 3), some of which occur on every *t*, while others alternate and only occur on every even *t* (such as *t* = 2, *t* = 4 and so on), or every odd *t* (such as *t* = 1, *t* = 3). It also includes functions that update and track the point scores achieved.

At the beginning of every iteration, the agents are each assigned a random x-coordinate and a random y-coordinate. Within the iteration, at the beginning of every *t*, both agents move randomly either 0 or 1 squares, and in any direction (figure 2). This means that on some *t*, the agent will move 0 squares and remain in their current location. To prevent them from moving off of the patch, they are forced to move backwards if they have reached the edge of the patch (i.e. an x or y-coordinate that equals 20). If both agents are on the same square (an encounter), the iteration loop breaks, the current iteration is terminated early, and the next iteration begins (where both agents are again placed on a random location on the patch). Details of the encounter logic are outlined in more detail under Submodels *Encounters*.

**Figure 2:**
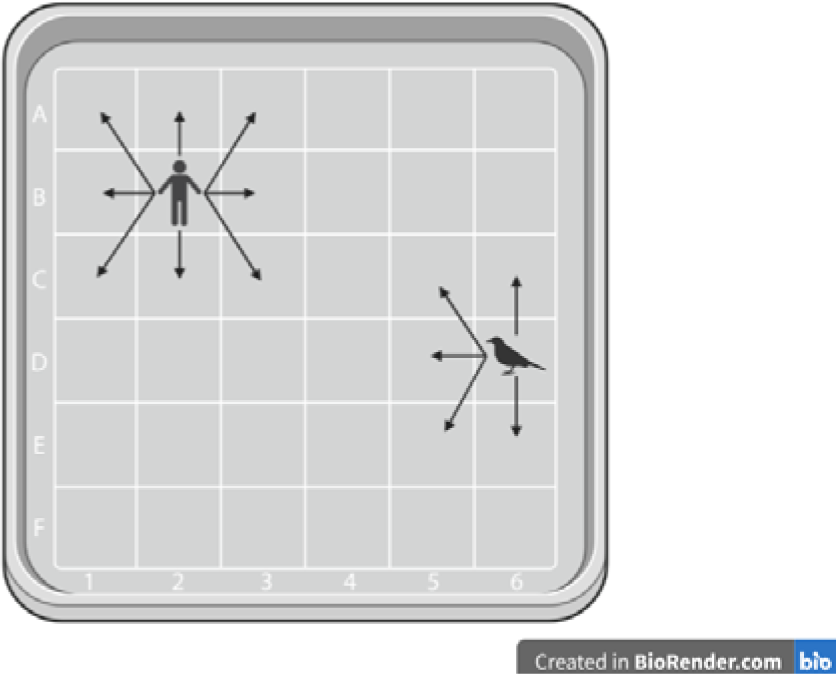
Illustration of the movement options for both agents. Each randomly moves 0-1 square in any direction, but they cannot move off the patch.

On every even *t*, the crow executes its foraging behaviour and then its eavesdropping behaviour. On every even *t*, the short-range scanning is executed. Long-range scanning is also executed on every even *t* unless the foraging behaviour has already detected the location of the human. The monitoring behaviour is executed on every odd *t* only if either foraging or long-range scanning have detected the location of the human in the current iteration. Details of these behaviours are outlined under Submodels *Foraging*, *Eavesdropping*, *Long-Range Scanning*, Short*-Range Scanning*, and *Monitoring*.

These behaviours alternate because a crow cannot simultaneously look around and forage on the ground, as they lower their head towards the ground to pick up food with their beak, and then periodically lift their head again to scan the environment (personal observations). Foraging and scanning therefore cannot occur within the same *t*. Eavesdropping is not restricted by head direction, so can be executed while foraging. It could theoretically also be executed while scanning, but as visual information is more reliable, visual information should be preferred over acoustic information. Speech eavesdropping has a high rate of false negatives where a human is present and not speaking, and it is less precise than visual scanning with regards to the distance of the human.

Finally, point scores are updated based on the events of the current *t*. These are outlined in more detail in the Submodels sections of the corresponding behaviours.

### 2.4 Design Concepts

#### Basic principles

At its core, the model addresses trade-off decisions between vigilance and foraging. An individual cannot just allocate all its time on a patch to foraging alone, as the risk of predation would be too high. It also cannot just allocate all its time to vigilance alone, as it would die from starvation. How much time must be allocated to vigilance, and how much can be spared for foraging, depends in part on the efficiency of the individual’s vigilance behaviours – the more efficiently and reliably it can detect predators, the more time is available for foraging as shown theoretically by Brown (1999). The present model extends this approach by examining the effect of introducing an additional vigilance behaviour in the context of urban habitats, where individual carrion crows can listen for human speech as an indicator of human presence (Schalz, 2023).

#### Emergence

The primary outcomes of the model are the difference in the crow’s survival rates with and without eavesdropping. Eavesdropping in addition to visual scanning is redundant under some environmental conditions, but beneficial under others. Exact threshold values are imposed by the point system of the model, but the existence of these thresholds emerges from the fundamental relationship between predation risk and vigilance efficiency (Brown, 1999).

#### Adaptation

The crow’s adaptive behaviour in this model is the relocation onto a different area of the patch once it has collected information showing that the human is nearby. Abandoning the patch entirely when the human is detected would result in the crow surviving only a handful of iterations before it died from starvation, as it could not collect enough points from foraging to compensate for its regular energy expenditure (based on an earlier pilot version of this model). Such a behaviour would prevent real crows to foraging in urban areas, as there are virtually no green spaces free from human presence. In the final version of the model, the crow instead relocates to a new square on the patch and so attempts to distance itself from the human without losing the foraging opportunity on the patch.

#### Objectives

The crow’s primary objective in the model is to increase its point score, which broadly represents its health and energy intake. Through its vigilance behaviours and subsequent relocation to a different area on the patch, the crow maximizes its point intake through foraging as often as possible, and minimizes its point losses by avoiding encounters with the human.

#### Predictions

The crow does not follow explicit predictions, but instead follows the implicit prediction that a current close proximity to the human increases the risk of future encounters, which then motivates the relocation onto a different area of the patch if the human is detected close by.

#### Sensing

At the beginning of each iteration, the crow has no information about the presence or location of the human on the patch. It collects this information through its vigilance behaviours; short-range scanning, long-range scanning, and eavesdropping. Once detected, it continuously updates this information through the monitoring behaviour, which is only reset if the crow relocates to a different area of the patch.

#### Interaction

When both agents are on the same square, the human injures the crow, resulting in a loss of points for the crow. The crow then abandons the patch in response, losing points through the energy expenditure of flight. It does not return to the patch until the next iteration starts. This flight response is based on previous experiments on flight initiation distances in wild birds, including American crows and hooded crows, taking flight when they detect a human below a certain distance threshold (Clucas & Marzluff, 2012). Carrion crows in close proximity to an audio speaker respond with flight behaviours and cessation of foraging when speech recordings are played (Schalz, 2023).

#### Stochasticity

The initial location of both agents is allocated randomly at the start of the iteration. Their movement on the patch is in a random direction within a set limit of steps. The points which the crow can gain from foraging or lose through encounters and flight are random within a set range of possible whole number values. The success of the eavesdropping behaviour has a set likelihood below 100%, and in runs of the model with reduced patch visibility, the success of visual scanning also has a set likelihood below 100%.

#### Observation

Throughout the model, a log file tracks the increase and decrease of the point scores. A second log file tracks the reason for terminating each iteration, i.e. whether it was fully executed (all 100 *t* completed) or whether it was terminated early after the crow had abandoned the patch. Both log files can be exported into csv files to see not only the final point score, but also how the score developed throughout the course of the model and how often encounters occurred.

Learning and collectives were not implemented.

### 2.5 Initialization

At the beginning of the first iteration, the starting point score for the crow is set. In subsequent iterations, the score is carried over from the previous iteration. While the human also has to have a point score for the point function to be executed correctly, it is not relevant to the model and set to a very high number so that it will not reach 0. All logs and the iteration counter are reset at the beginning of the model.

At the start of each iteration, the counter for time steps *t* is reset to 1, and the x and y-coordinates of both agents is randomly assigned. At the start of an iteration, the crow does not have any information about the human, so these variables are set to FALSE.

### 2.6 Input Data

The model does not use input data to represent time-varying processes, it is fully self-contained from its internal parameter values set in the script.

### 2.7 Submodels

#### Foraging

On every even *t* (for instance *t* = 2, *t* = 4 etc), if the crow has not yet reached it’s maximum score of 25,000 points, it engages in the foraging behaviour and gains randomly between 10 and 65 points. The lower and upper point limits correspond to the caloric value of an earthworm (Finke, 2002) and a piece of Dorito (Doritos UK, n.d.) respectively, two food items that a crow might find in an urban public park. Food is eaten immediately and cannot be taken away or cached for later, so the points are added to the crow’s overall score immediately. Crows cannot store an infinite amount of energy in their bodies, so the model crow will not take in any additional points if its current point score exceeds a maximum capacity of 25,000 points (which is 4,000 points above its starting point score).

#### Eavesdropping

On every even *t*, if the crow has not already detected the human’s location, it engages in the eavesdropping behaviour. This behaviour does not occur on odd *t*’s during visual scans (see below), as visual information would be more accurate and should be preferred over acoustic information when available. The eavesdropping behaviour therefore provides low-accuracy information during foraging, when no other information is available to the crow.

Humans only speak for approximately 20% of the day (Janusik & Wolvin, 2009). The crow therefore only has a 20% chance of detecting the human through eavesdropping (figure 4). This is represented in the model script by a one out of five chance that the behaviour is executed. When it is executed, the human’s location is always detected. This means that the crow can hear the human occasionally, but will mostly receive false negatives as the human is present but not speaking.

**Figure 3:**
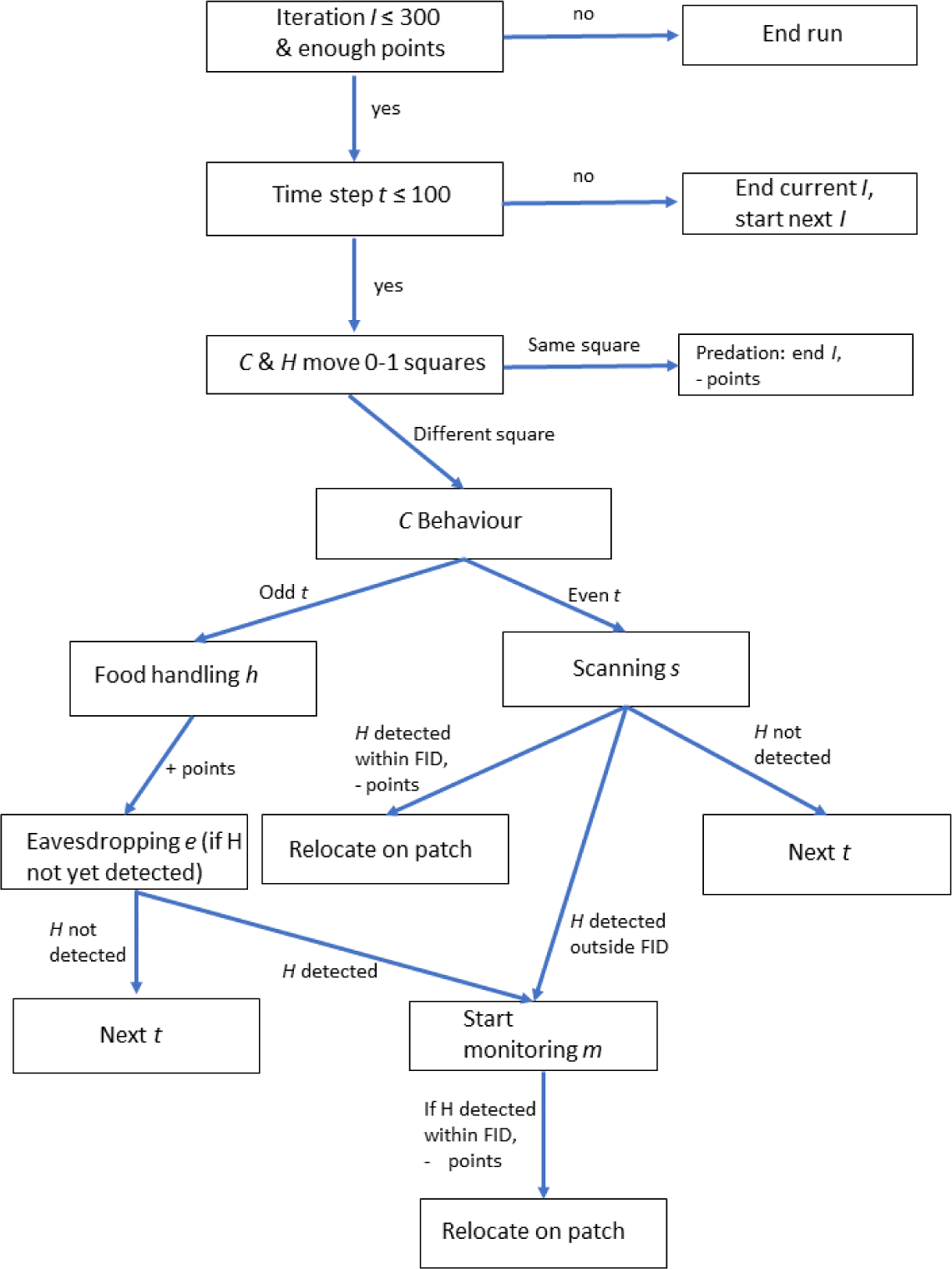
Flowchart of the model logic, showing the order of behaviours and steps of the model. Iteration is shortened to I, crow to C, and human to H.

**Figure 4:**
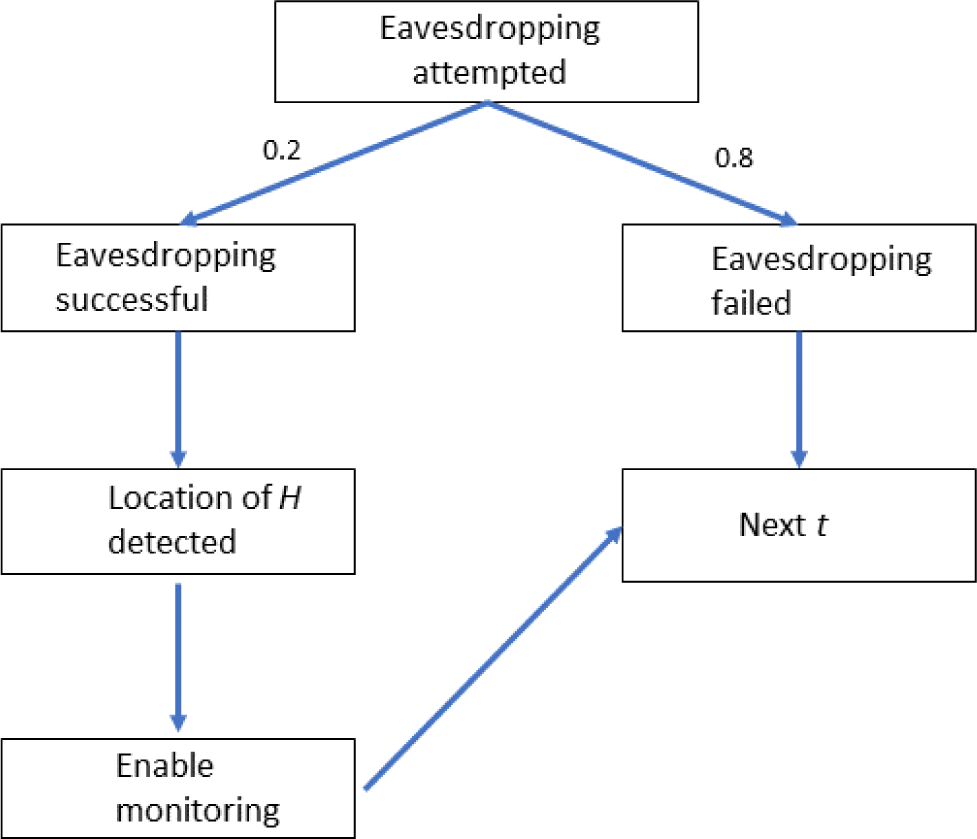
Flowchart of the eavesdropping behaviour logic. When eavesdropping is attempted, there is a 20% chance that it will be successful, and an 80% chance that it will fail. If successful, the location of the human is detected, and that location is monitored from that *t* onwards.

#### Long-Range Scanning

On every odd *t* (for instance *t* = 1 or *t* = 3), if the crow has not already detected the human’s location, long-range scanning is executed. This behaviour samples a random row of squares on the patch (for example all squares with an x-coordinate of 3, see figure 5). Scanning one row at a time is meant to imitate the range restrictions of visual scanning (only one direction at a time), but allowing for multiple squares in one line of sight to be seen. The same squares could be randomly sampled multiple times, as the human’s movements mean that a previously empty square could later be occupied. Squares outside the patch cannot be sampled.

**Figure 5:**
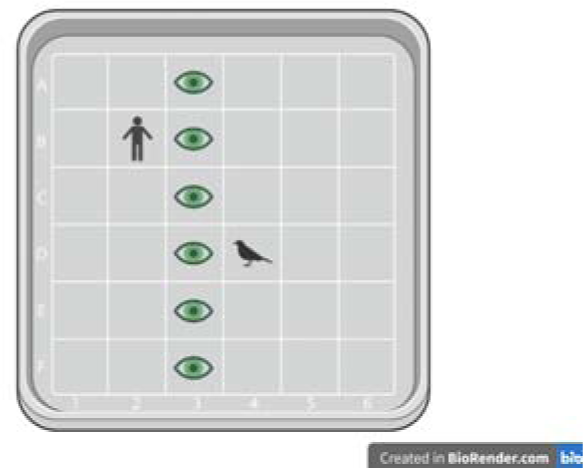
Example illustrating the long-range scanning. The crow samples an entire randomly selected row of squares (marked with green eyes), in this case all squares with the x-coordinate 3. The human is not located on any of these squares, so would not be detected in this scan.

#### Short-Range Scanning

On every odd *t*, the crow executes its short-distance scan and randomly samples two squares within a two square radius of its current location (figure 6). Sampling two random squares that may not necessarily be adjacent to each other (e.g. C4 and D2) deviates from the natural visual range of a crow, but was chosen for the sake of model simplicity. As the human moves randomly as well, they may walk to any square with equal likelihood, so the crow scanning randomly has the same odds of spotting the human as it would if it would only scan pairs of adjacent squares. This deviation therefore does not impact the model outcome. Short-distance scans are particularly useful to avoid an encounter before eavesdropping or long-range scanning have detected the human’s location, as this behaviour focuses on the crow’s immediate surroundings. As with long-range scanning, the same squares could be sampled multiple times, but squares outside the patch cannot be sampled.

**Figure 6:**
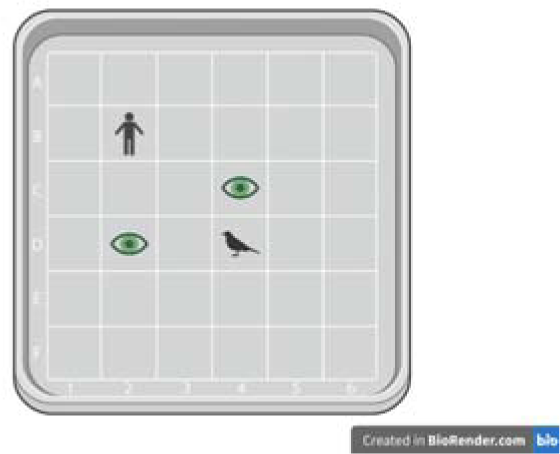
Example illustration of the short-distance scanning behaviour. Out of all squares within a two square radius of the crow, it randomly samples two squares: in this case, C4 and D2, marked with green eyes. The human is not located on either of these squares, so would not be detected in this scan.

Foraging and scanning alternate (instead of occurring in a randomized order) for two reasons: wild crows roughly alternate these behaviours rather than engaging in them completely randomly (personal observations). Additionally, a randomized order of these behaviours would influence the model outcome beyond the effect of the eavesdropping behaviour, which is the focus of this paper (e.g. a human encounter occurs not because of failed eavesdropping, but because of the foraging behaviour having been selected randomly for the past five *t*).

The crow engages in both long-range and short-range scans for conceptual reasons: short-range scans trigger an immediate need to take flight, as they only include squares within the crow’s flight initiation distance (FID) range, i.e. the human is already too close to the crow. Long-range scans provide information on the presence of humans on the overall patch, but as the human can be far away no immediate need for flight is present.

#### Monitoring

If the human’s location is detected through the visual scanning or the eavesdropping behaviour, the crow begins to track the human’s location on every odd *t*. This replaces the visual scanning behaviours, and also disables the eavesdropping behaviour. In other words, it continues foraging on an even *t*, then gets an update on the human’s position on the following odd *t*, and then forages again on the next even *t*. This means that encounters would only occur if the human reaches the crow before being detected by any of the vigilance methods. The patch is only abandoned if the crow and the human move onto the same square.

Monitoring the human’s location is preferable to simply abandoning the patch, as it allows the crow to continue foraging while the human is still at a safe distance. If the monitoring behaviour detects the human within less than 3 squares of the crow, the crow relocates to a new area of the patch (figure 7). This represents the “hopping away” behaviour displayed by urban crows. Relocating costs 2 points in energy loss, plus an additional 10 - 65 penalty (representing the lost value of dropping the food of the current square and losing out on this one foraging opportunity). After relocating, the monitoring behaviour resets, and the crow has to newly discover the location of the human before monitoring can be used again.

**Figure 7:**
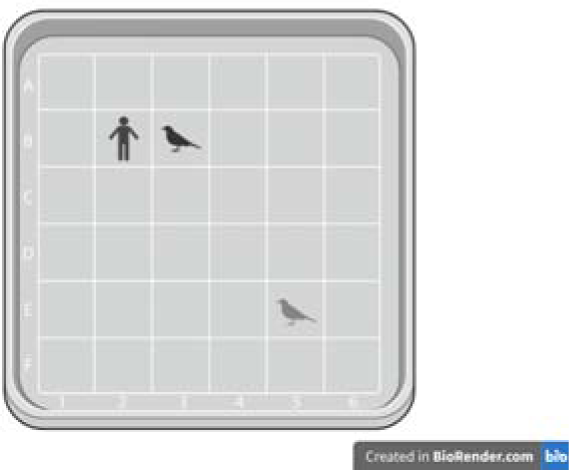
Illustration of the relocating behaviour. Once the crow has detected the human within 2 squares of its own location (in this example on squares B2 and B3 respectively), it relocates to a random position on the patch (in this example E5).

#### Encounters

If the human and the crow are on the same square at the same time, the crow will first lose points due to injury (randomly between −1000 and −3,500 points). It will then flee from the patch, which costs another 2 points, representing the energy expenditure of a 5s flight (an arbitrary flight duration from the patch to a hypothetical perching spot off the patch), at 0.00765kcal/s of flight (see Zach, 2008). As the crow has abandoned the patch, the iteration ends immediately and the next iteration starts. An early termination of the current iteration also means that the crow loses out on any points it may have gained in the remaining *t* of the iteration. This is a third, although indirect, cost of the encounter.

### 2.8 Simulation Experiments

To ensure all parameters of the model interact as expected, I completed 20 test runs (i.e. running the full model with its 300 iteration 20 separate times) at each design stage of the model. In the final version outlined in this section, the crow had near-perfect survival rates (median final point score was 24,978, which is only 22 points below the highest possible score; SD = 1,723 points). The crow successfully detects and evades the human most of the time, and the lost points from rare encounters can be compensated in future iterations.

Once this final version was achieved, I systematically altered the environmental parameters of the model. The aim was to test under which environmental conditions eavesdropping on speech provides a benefit to the crow, i.e. under which conditions the crow survives more often and collects more points when eavesdropping is available. To do this, I separately altered the patch size, the share of the patch that can be scanned due to visual obstacles, as well as the count of humans present on the patch. Each variation was tested individually (i.e. only one parameter was modified at a time). In each variation, I also modified the available vigilance behaviours, to compare the effect they have on the crow’s performance (table 2).

**Table 1:** Iterations and time units in the model.

| Time | Corresponds to |
| --- | --- |
| 1 time unit $t$ | smallest time segment in the model |
| 100 time units $t$ | 1 iteration |
| 1 iteration | 30 minutes |
| 15 iterations | 1 day |
| 300 iterations | 20 days, full length of the model |

**Table 2:** Combinations of environmental settings and available vigilance behaviours for each environmental parameter altered. Each pair was tested individually, e.g. 20 runs of the model with a 10 x 10 square and eavesdropping enabled, then 20 runs of the model with a 10 x 10 square and eavesdropping disabled, and so on.

| **Environmental Variable** | **Versions of the Environment** | **Available Vigilance Behaviours** |
| --- | --- | --- |
| Patch Size | 10 x 10 squares<br>13 x 13 squares<br>14 x 14 squares<br>15 x 15 squares<br>17 x 17 squares<br>20 x 20 squares | Eavesdropping + visual scanning<br>Visual scanning only<br>None |
| Patch Visibility<br>(patch size:<br>17 x 17 squares) | 100%<br>75%<br>50%<br>20%<br>5%<br>0% | Eavesdropping + visual scanning<br>Visual scanning only |
| Count of Humans | 1 human, 20 x 20 squares<br>1 human, 22 x 22 squares<br>1 human, 25 x 25 squares | Eavesdropping + visual scanning<br>Visual scanning only<br>None |
|  | 1 human, 30 x 30 squares |  |
|  | 1 human, 40 x 40 squares |  |
|  | 2 humans, 20 x 20 squares |  |
|  | 2 humans, 22 x 22 squares |  |
|  | 2 humans, 25 x 25 squares |  |
|  | 2 humans, 30 x 30 squares |  |
|  | 2 humans, 40 x 40 squares |  |

Each combination of environmental variation and available vigilance behaviours was tested in 20 separate runs of the model (e.g. 20 runs with eavesdropping, and 20 runs without eavesdropping enabled, for each version of the environment). This number of runs per combination was predetermined during the model exploration stage.

## 3. Results

### 3.1 Variation in Patch Size

For crows foraging in real habitats, patch sizes will vary and may at times be very small. The smaller the patch size, the more likely an encounter with the human, and the less time for the crow to detect them. To survive all iterations without any vigilance behaviours (eavesdropping and scanning disabled) required a minimum patch size of 20×20, whereas the minimum size with visual scanning enabled was 17×17. When full vigilance monitoring (eavesdropping and visual scanning) was used, the minimum size decreased further to 15×15 (figure 8). On smaller patches, versions of the model with more monitoring behaviours performed better than those with fewer detection behaviours, because the human needed to be detected quicker.

**Figure 8:**
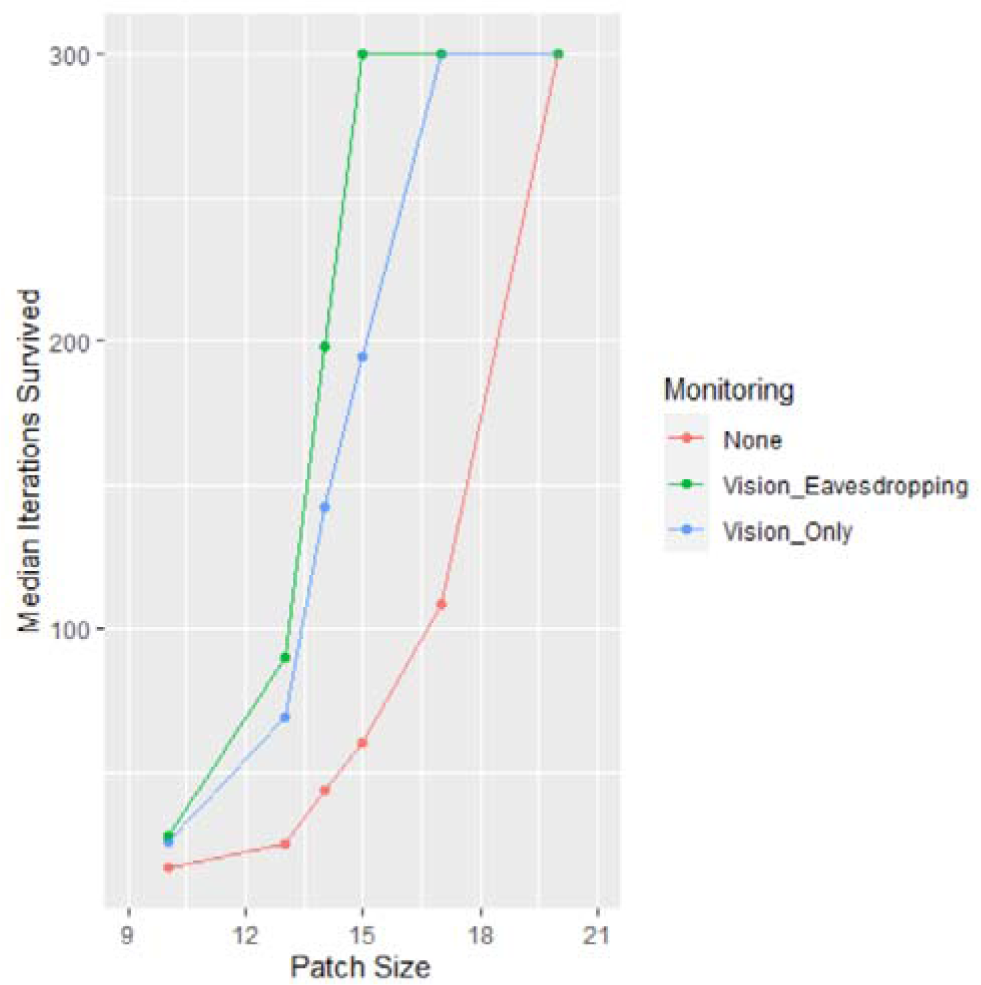
Median survived iterations across 20 runs per setting, each with a variation of patch size (10×10, 13×13, 14×14, 15×15, 17×17, 20×20) and monitoring abilities (both visual long-range scanning and eavesdropping enabled, only visual scanning enabled, and neither one enabled). Created with the R package ggplot2 (Wickham, 2016).

### 3.2 Variation in Visual Obstacles

Some patches may have objects like trees, shrubs, buildings or cars. These visual obstacles reduce the accuracy of visual scanning, as only some areas of the patch can be covered with this vigilance method (figure 9). This was modelled implicitly by reducing the scanning accuracy from (100% down to 75%, 50%, 25%, 5% and 0%), reflecting that a certain percentage of the square could not be scanned visually.

**Figure 9:**
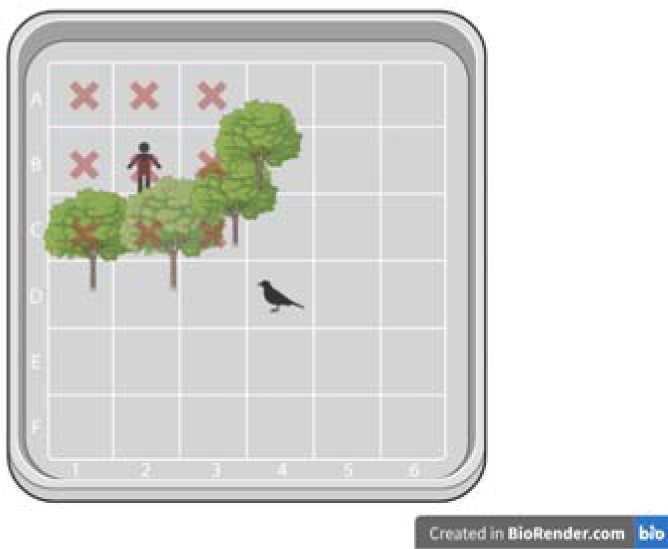
Some squares on the patch are hidden behind trees as visual obstacles, which means the crow cannot visually scan squares A1-A3, B1-B3, or C1-C3, including the square of the human. These obstacles reduce the accuracy of its visual scan below 100%.

When eavesdropping was enabled, survival rates were perfect even if long-range scanning was impossible (i.e. in runs where 0% of the patch was visible). When eavesdropping was disabled and only long-range scanning was enabled, survival rates dropped below 300 iterations when visual scanning accuracy was less than 100% (figure 10). Eavesdropping is therefore beneficial when obstacles reduce the crow’s visual field and prevent it from seeing the human in all instances, on a patch that is small enough for encounters to occur regularly.

**Figure 10:**
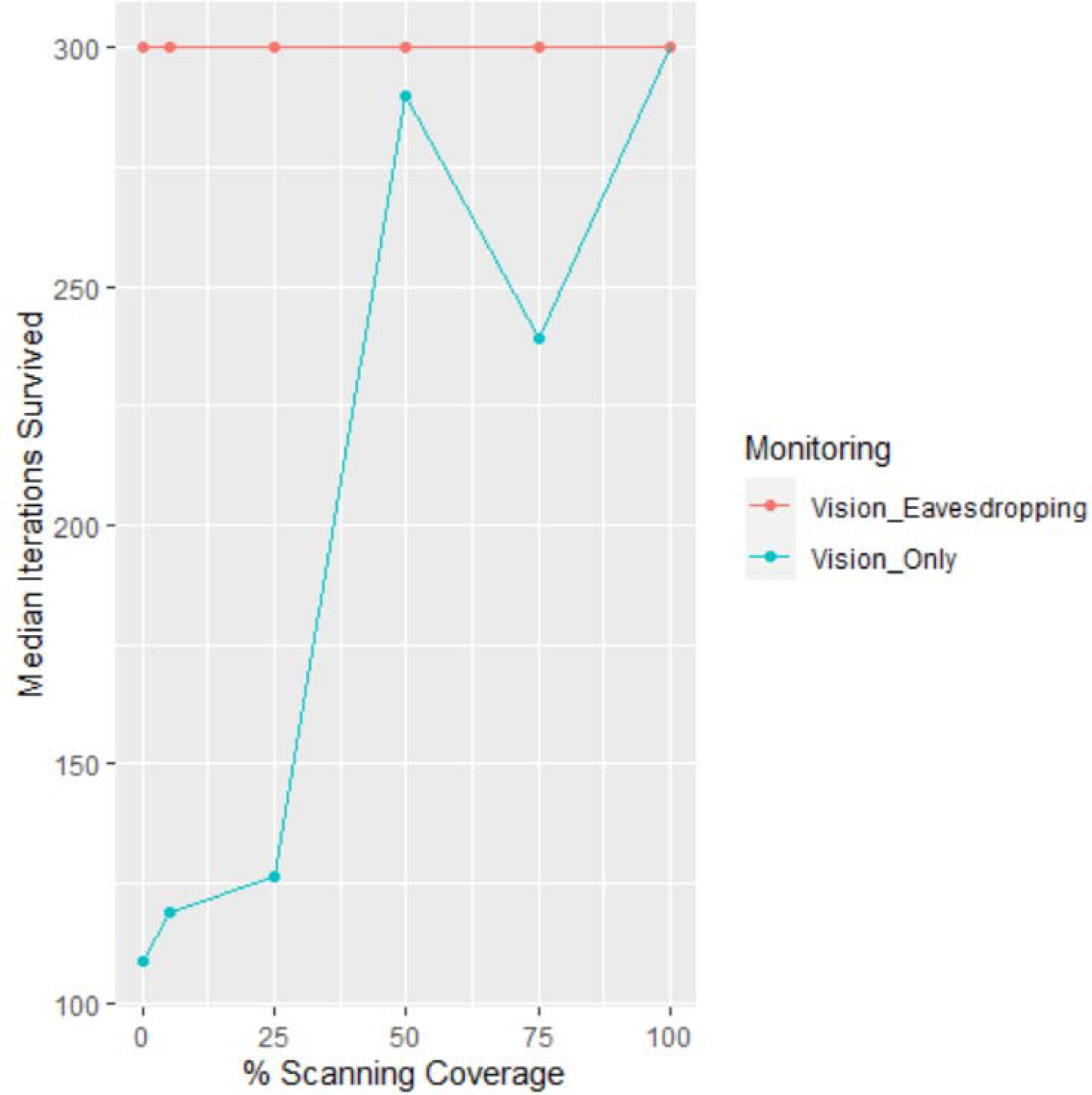
Median iterations survived with eavesdropping enabled (red) or disabled (blue), with variable scanning coverage of the patch to represent the percentage of patch area that is scannable and not covered by obstacles. Created with the R package ggplot2 (Wickham, 2016).

### 3.3 Variation in Human Presence

Real crows will experience different human densities and may need to avoid more than one human on a given patch. I modelled the survival rates of the crow with an additional human placed on the 20×20 patch. As before, I conducted 20 runs for each combination of human density and vigilance method. Both humans move independently of each other but follow the same walking style, and the crow responds to both in the same way as previously specified.

While the crow had perfect survival odds with one human on a 20×20 patch, the median number of survived iterations with two humans and full monitoring (eavesdropping and long-range scanning) decreased to 96 iterations. The crow only survived all 300 iterations when the patch size was increased to 22×22 squares. As in Brown’s (1999) vigilance model, survival chances are proportionate to the predation risk (i.e. the risk of encountering one of the humans), as well as the employed vigilance levels (in this case either none, visual scanning only, visual scanning and eavesdropping).

When only long-range scanning was enabled and eavesdropping was disabled, the minimum patch size for perfect survival with two humans was 25×25. With both methods disabled, a minimum patch size of 30×30 was needed. As the human density increases, higher vigilance efficiency is needed and eavesdropping becomes increasingly beneficial. But even with the full set of vigilance behaviours enabled, the patch still needed to be 2.15 times larger for the crow to have the same survival rates as it did with only one human (figure 11 and table 3).

**Figure 11:**
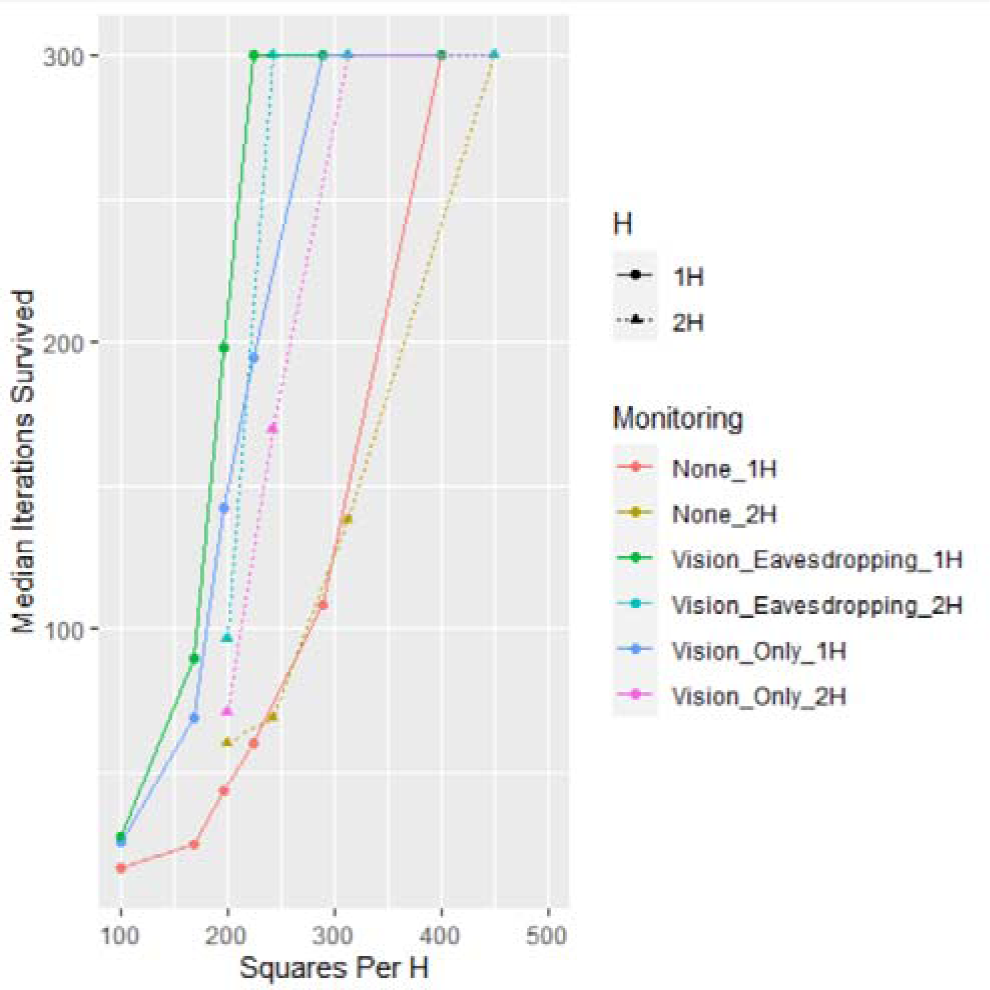
Median survived iterations across 20 runs per setting across different densities (squares per human H), with 1 H (solid line, round points) or 2 H (dotted line, triangle points) and different monitoring abilities (both visual long-range scanning and eavesdropping enabled, only visual scanning enabled, and neither one enabled). Created with the R package ggplot2 (Wickham, 2016).

**Table 3:** Number of squares (i.e. a 20×20 patch has 400 squares) on the smallest possible patch with perfect survival for each monitoring setting and either one or two humans. Patch Size Increase Factor is the relative increase in patch size from 1 to 2 humans.

| Monitoring | Minimum Squares 1 Human | Minimum Squares 2 Humans | Patch Size Increase Factor |
| --- | --- | --- | --- |
| None | 400 | 900 | 2.25 |
| Vision Only | 289 | 625 | 2.16 |
| Vision and Eavesdropping | 225 | 484 | 2.15 |

## 4. Discussion

With this model, I have aimed to show under which environmental conditions eavesdropping on human speech may provide a benefit to carrion crows, in the form of increased survival for individuals engaging in this behaviour. To do this, I have used an agent-based model to create a baseline set-up of a crow foraging on a patch while trying to avoid encountering a human. Parameters were set to be as realistic as possible, such as aligning the point system with the calory needs and energy expenditures of real crows. Once the model was fully set up, I systematically varied the environmental conditions of the patch to see when eavesdropping would result in higher survival rates than visual scanning on its own.

I found that eavesdropping results in increased survival rates when human density is increased (either because the patch is small, or because several humans are present). Additionally, eavesdropping led to higher survival rates when parts of the patch could not be visually scanned due to obstacles. A large, open field would have the lowest risk and should not need eavesdropping, as visual scanning would be fully sufficient. Small, crowded patches or patches with obstacles on the other hand have a higher risk of an encounter occurring before the crow can visually detect the human, so acoustic monitoring via speech eavesdropping provides a benefit there.

There are some limitations to the conclusions that can be drawn from the model. A theoretical model cannot include all variables of real-world scenarios, and there may be other factors influencing the survival rates and eavesdropping benefits for wild crows. The present model did not include factors such as the crow’s internal state, its breeding activities, or the presence of conspecifics. A hungry crow should take higher risks during foraging to prevent starvation, whereas a crow that has had sufficient food can afford to avoid riskier patches entirely. Likewise, a crow foraging to feed its chicks may need to accept higher risks in order to collect enough food. If conspecifics are present, alarm calls would provide the focal individual additional information beyond its own vigilance efforts.

Another factor that was not included in the model is previous experience with humans. Having been repelled or chased by humans in the past (Clucas & Marzluff, 2012; Schalz, 2021) should influence the crow’s overall risk perception of humans; a crow that has only been ignored or fed by humans should tolerate a higher risk than a crow that has experienced chasing.

The model assumed a 20% likelihood of any eavesdropping attempt to be successful, based on the share of time humans generally speak in a given day. However, most versions of the model were set up with only one human present. While crows could encounter a single human talking on the phone or talking to themselves, I would expect this to be relatively rare. When two humans are on the patch together, on the other hand, speech rates should be higher as they would be more likely to talk to each other. This suggests that the success of speech eavesdropping may vary between different types of patches, and may for instance be more successful in parks where people are likely to go on walks with friends and family.

Finally, the model was designed under the assumption that eavesdropping does provide a benefit, given that this behaviour has been shown in crows of different species (Schalz, 2023), and different sites (Schalz & Izawa, 2020). While it was possible for the model to show a negative effect of eavesdropping on survival, and while it did show a lack of effect in some scenarios, there is an inherent bias in the design towards finding positive rather than negative effects.

Overall, the model suggests that crows foraging on patches that have a high human density, or have visual obstacles such as trees would benefit from eavesdropping on speech as an additional vigilance behaviour. Future empirical research could examine whether crows respond more to playback of speech when foraging on these types of patches compared to lower-risk alternatives. Entering the measurements and conditions of existing patches in the model may help to select experimental patches to compare. Long-term studies on the survival rates of crows relative to their individual responsiveness to speech playback would also provide valuable insights. Lastly, while the parameters were defined with carrion crows in mind, this model may also be applicable to other avian species. A cross-species comparison of speech eavesdropping on different patch types may motivate further research into this currently under-investigated behaviour.

## Acknowledgements

I am very grateful to Tom Dickins and Martijn Timmermans for their support during the conceptualisation of the model, as well as their feedback on an earlier draft of this manuscript. Likewise, I thank the anonymous reviewers who have provided valuable feedback so far, which has been incorporated into this latest version of the manuscript. I also thank Brad Duthie for making his R AGM script and tutorial open-access, which was extremely helpful during the first stages of writing my own model script.

## Data Availability

R scripts and data generated in the model can be accessed freely via: https://doi.org/10.6084/m9.figshare.c.6651116.v3

## Conflict of Interest

The author declares no conflict of interest.

## Notes

https://doi.org/10.6084/m9.figshare.c.6651116.v3

### Competing Interest Statement

The authors have declared no competing interest.

### Summary of Updates

Updates throughout to address comments and suggestions from reviewers.

